# Defining Cardiac Nerve Architecture During Development, Disease, and Regeneration

**DOI:** 10.1101/2022.12.31.522405

**Authors:** Rebecca J. Salamon, Poorva Halbe, William Kasberg, Jiyoung Bae, Anjon Audhya, Ahmed I. Mahmoud

**Affiliations:** Department of Cell and Regenerative Biology, University of Wisconsin School of Medicine and Public Health, Madison, WI 53705, United States; Department of Biomolecular Chemistry, University of Wisconsin School of Medicine and Public Health, Madison, WI 53705, United States; Department of Nutritional Sciences, Oklahoma State University, Stillwater, OK 74078, United States

**Keywords:** cardiac nerves, cardiac regeneration, myocardial infarction, coronary vasculature, choline acetyltransferase, parasympathetic innervation, connexin 40

## Abstract

Cardiac nerves regulate neonatal mouse heart regeneration and are susceptible to pathological remodeling following adult injury. Understanding cardiac nerve remodeling can lead to new strategies to promote cardiac repair. Our current understanding of cardiac nerve architecture has been limited to two-dimensional analysis. Here, we use genetic models, whole-mount imaging, and three-dimensional modeling tools to define cardiac nerve architecture and neurovascular association during development, disease, and regeneration. Our results demonstrate that cardiac nerves sequentially associate with coronary veins and arteries during development. Remarkably, our results reveal that parasympathetic nerves densely innervate the ventricles. Furthermore, parasympathetic and sympathetic nerves develop synchronously and are intertwined throughout the ventricles. Importantly, the regenerating myocardium reestablishes physiological innervation, in stark contrast to the non-regenerating heart. Mechanistically, reinnervation during regeneration is dependent on collateral artery formation. Our results reveal how defining cardiac nerve remodeling during homeostasis, disease, and regeneration can identify new therapies for cardiac disease.

## INTRODUCTION

The cardiac nervous system is a pillar of cardiac physiology, regulating conduction and contractility that is balanced by two opposing branches of the sympathetic and parasympathetic input^1^. Cardiac nerves localize to specialized nodes, the heart’s pacemaker cells, where signals for contraction are propagated throughout the atria and into the ventricles^2,3^. Recent studies led to new insights on the patterning and function of the intrinsic cardiac nervous system, which underscores the importance of dissecting the role of neurocardiology in cardiac health and disease^4, 5^.

The influence of cardiac innervation expands beyond their well-known autonomic role in the heart. Specifically, sympathetic nerves have been demonstrated to extensively innervate the ventricles^6, 7^. These ventricular nerve networks mediate communication across a variety of cell types, including myocytes and blood vessels^8, 9^. Neural connections are established during late embryonic and early postnatal development by a co-maturation system^10, 11^ where innervation patterning and maturation relies on signals from a variety of cardiac tissues^12, 13^. Interestingly, sympathetic nerves have been demonstrated to mediate the postnatal transition of cardiomyocytes from hyperplasia to hypertrophy^14^ and cardiomyocyte size^15^.

In contrast, parasympathetic innervation has been less studied for its interactions within the cardiac ventricles, due to the widely-held belief that parasympathetic nerves do not significantly contribute to ventricular innervation^16^. This misconception has recently been challenged by evidence of parasympathetic nerve function within the ventricles, including a high density of cells expressing muscarinic receptors^17, 18^, protective functions against ventricular arrythmias^19^, effects on contractility influence^20^, and distribution of parasympathetic nerve fibers^21–24^ Yet, the anatomy, distribution, and cell-cell interactions of ventricular parasympathetic innervation remains unclear.

Autonomic nerve remodeling and dysfunction contributes to the pathogenesis of cardiovascular disease^10, 25^. The heterogeneity of nerve remodeling, including regions of denervation and sympathetic hyperinnervation, has been demonstrated to contribute to ventricular arrhythmias and sudden cardiac death^25–29^. Interestingly, nerves play an important role in promoting repair and regeneration across a variety of organs and species^30^. Importantly, cholinergic nerve function regulates cardiomyocyte proliferation and regeneration of the neonatal mouse heart^31^. Remarkably, this regenerative response is accompanied with restoration of autonomic functions, suggesting physiological innervation of the regenerating mammalian heart^32^. However, how cardiac nerve patterning and distribution compares to the uninjured or diseased heart has not been explored.

The broadened role of the intrinsic cardiac nervous system in regulating heart development, homeostasis, and regeneration highlights the importance of defining the patterns of cardiac innervation as a necessary step to promote physiological reinnervation in many cardiomyopathies. Gaining insights into the patterns of cardiac innervation and nerve density during development is necessary to understand the normal circuitry of cardiac innervation. Our current understanding of cardiac nerve patterns is largely based on two-dimensional (2D) histological analysis, where fixed tissues are labeled with different markers for visualization of neurons. This approach is limited as it provides no information regarding their spatiotemporal distribution. Additionally, histological analysis cannot reveal information about the developmental distribution of neurons in the heart.

Thus, comprehensive lineage tracing of individual cardiac nerves while visualizing them in a three-dimensional (3D) manner is necessary to dissect the development, maturation, and lineage commitment of the cardiac autonomic nervous system.

Here we generate and analyze the first 3D neurovascular map of the mature, intact murine heart ventricles. We reconstruct nerve patterning with high-spatial accuracy by employing comprehensive lineage tracing, whole-mount immunostaining, confocal microscopy, and 3D reconstruction. We use this system to further explore neurovascular development and distribution of the sympathetic and parasympathetic subpopulations. We demonstrate that parasympathetic nerves extensively innervate the heart during development and in the postnatal heart together with sympathetic nerves, highlighting that parasympathetic nerves contribute significantly to ventricular innervation. Furthermore, we demonstrate physiological nerve remodeling during regeneration, which is distinct from pathological innervation of the non-regenerating heart. Mechanistically, we demonstrate that physiological reinnervation during regeneration is dependent on collateral artery formation. Together, our findings reconstruct cardiac nerve architecture and remodeling during development, disease, and regeneration.

## METHODS

### Mice

All animal experimental procedures were approved by the Institutional Animal Care and Use Committee of the University of Wisconsin-Madison. All experiments were performed on age and sex matched mice, with an equal ratio of male to female mice. Mouse lines used in this study are: *ChATCre* (The Jackson Laboratory, Stock# 006410), *Rosa26^tdTomato^* (The Jackson laboratory, Stock# 007905), *Cx40CreER* (^33^, *Cxcr4^fl/fl^* (The Jackson laboratory, Stock# 008767).

### Tamoxifen Administration

Tamoxifen was prepared at 100 mg/ml, dissolved in a 9:1 solution of corn oil to 100% ethanol, and incubated at 37°C overnight with rotation. Solution was vortex as needed. Tamoxifen stock was kept at 4°C for up to one month and incubated at 37°C overnight before use. For *Cx40CreER* postnatal Cre induction, tamoxifen (1mg per pup) was administered by a subcutaneous (SubQ) injection directly to pup at P4^34^ For *Cx40CreER* embryonic Cre induction, tamoxifen (0.1mg/g BW) was administered to the intraperitoneal (IP) cavity of the pregnant dam 24 hours before harvest.

### Myocardial infarction surgery

Myocardial Infarction surgery (MI) was performed at P1 or P7, as described^35^. Pups were separated from dam and placed into a new cage with bedding. Pup was anesthetized on ice for 3 minutes. Working under a dissecting scope, a small incision was made in the skin, area under the skin was loosened, the 4^th^ intercostal muscle was located, and incision was made, and the heart was gently guided out of the chest cavity with blunt forceps. The LAD was located at ligated with a 6-0 prolene suture by a surgeon’s knot followed by a simple knot, and apical blanching was visualized. Heart was gently guided back into the chest cavity, ribs were sutured together, and skin was glued. Pup was placed into a heating pad until lively. Once surgery was complete, pups were rubbed with original bedding to transfer scent and placed back to dam. Hearts were collected at 7- or 21-days post-MI.

### Tissue clearing of intact postnatal hearts

Our passive CLARITY technique was performed with minimal modifications^36^ in hearts that required imaging of structures deep within the tissue (i.e. coronary arteries). Briefly, hearts were harvested, washed in PBS, and placed into a 2ml (embryonic to P7 hearts) or 4ml (P21+ hearts) glass vial with 4% PFA, incubating overnight at 4°C with rotation. Following, hearts were washed with PBS for 30min at room temperature (RT), repeated three times, and placed into a polymerization solution (4% acrylamide and 0.5% VA-044) overnight at 4°C. The next day, polymerization was activated with a 3-hour incubation at 37°C. Hearts were again washed at RT in PBS for 1 hour, repeated three times. Hearts were placed into clearing solution (8.0% w/v SDS, 1.25% w/v Boric acid, 0.5% w/v 1-thiioglycerol dissolved in purified H2O, pH 8.5) and incubated at 37°C, changing the solution every two days until tissue was fully cleared. Typically, P7 and P22 hearts were cleared for 5-7 or 14-16 days, respectively. In some tissues, a dark, green colored pigment persists even after clearing; regardless tissues were moved onto next steps, as it will be cleared out during washing. After clearing, tissues were washed in a conical tube with 10ml of PBS for 3 days, changing PBS solution 2-3 times daily.

### Whole-mount immunohistochemistry

For uncleared tissues, hearts were harvested, blood removed, placed into a 2ml vial (for embryonic to P7 hearts) or 4ml vial (older than P7 hearts) and fixed for 1 hour in 4% PFA at 4°C (for embryonic hearts) or RT (for postnatal hearts). Hearts were washed in PBS for 15 min, repeated 3 times.

Cleared and uncleared hearts underwent the same immunohistochemistry staining protocol described. Hearts were blocked in 20% blocking buffer (BB, matching serum from secondary host) made in PBS with 0.2% triton (PBST) for 1 hour at RT. Primary antibodies were diluted in 0.2% PSBT with 2% BB and at the following concentrations: rabbit RFP/tdTomato (Rockland, Cat# RL600-401-379) at [1:200], mouse Tuj1 (Sigma, Cat# T8453) at [1:200], rabbit Tuj1 at [1:500], sheep TH (Chemicon, Cat# AB1542) at [1:200], endomucin (Santa Cruz, Cat# SC65494) at [1:100]. Primary concentrations for hearts at P21 or older were doubled. Primary incubation was performed overnight at RT, then transferred to 4°C for an additional overnight incubation. Following, hearts were washed with PBS for 30min at RT, repeated three times. Secondary antibodies were diluted in 0.2% PBST + BB at [1:250], using the following Alexa Fluor antibodies from Invitrogen: 594 anti-rabbit (Cat# A32740), 488 anti-mouse (Cat# A32723), 488 anti-sheep (Cat# A-11015), 405 anti-rat (A48261). Secondary incubation was performed for 3 hours at RT then moved to 4°C overnight. When using a combination of mouse and rat primary antibodies, the staining was performed sequentially to limit cross-reaction of secondary antibodies^37^

### Microscopy and 3D reconstruction

Confocal imaging was performed on a Nikon Upright FN1 microscope equipped with high sensitivity GaAsP detectors. Hearts were placed into a 3D-printed well, filled with water, and positioned for anatomical view. Whole heart images were taken using a 4x air objective (0.2 NA), with 5 um spacing between z-planes, and tiles stitched with Nikon pic-stitching function, and scale bars shown as 1 mm. Higher magnification images were taken using a 40x water immersion objective (0.8 NA), with 1 um steps between z-planes, and scale bars shown as 100 um. Imaris microscopy image analysis software in filaments mode was used to segment hearts generate statistics. Representative images are shown as max intensity projections (MIP) and were edited using Adobe Lightroom and Photoshop for clarity. Since Endomucin (EMCN) stains veins and capillaries, EMCN signal in large-diameter veins was artificially highlighted in Photoshop. Confocal images were deconvolved using iterative classic maximum likelihood estimation in Huygens Profession before Imaris 3D reconstruction.

### Statistical Analysis

Data generated via Imaris, with 2-3 replicates averaged per sample and 3 samples per group. Graphs generated in GraphPad Prism 9, with individual points representing the average per sample. Groups compared by an ordinary two-way ANOVA, with uncorrected Fisher’s LSD, with a single pooled variance. Significance shown as n.s. (P > 0.05), * (P ≤0.05). **(P ≤0.01), ***(P ≤0.001), **** (P ≤0.0001).

## RESULTS

### Cardiac Nerves Sequentially Associate with Coronary Veins and Arteries During Development

The development of cardiac innervation throughout the ventricles of the embryonic heart is not completely defined. Our current understanding of cardiac nerve patterning of heart development is largely based on 2D analysis; however, this approach is insufficient to reconstruct the intricate nerve networks and cell-cell interactions. Specifically, although the coronary arteries are known to be innervated by sympathetic axons^13^, the developmental timeline and patterning of artery innervation is not well defined. Here, we sought to elucidate the dynamics between nerve-vein and nerve-artery association during embryonic heart development.

The main, large-diameter veins of the embryonic heart primarily reside on the posterior wall^13^, whereas the main coronary arteries are primarily located within the anterior wall^13, 38^. Therefore, we divided the architectural regions into posterior and anterior sides of the heart, including right and left ventricles (LV, RV) (**Figure 1**). Coronary arteries were visualized with lineage labeling using the inducible Connexin 40 Cre (*Cx40CreER*) and the *Rosa26^tdTomato^* reporter mice (**Figure 1**). Neurovascular patterning was identified with additional markers for all nerves using beta tubulin III (Tuj1), and veins were labeled with endomucin (EMCN) (**Figures 1A-C**).

**Figure 1.**
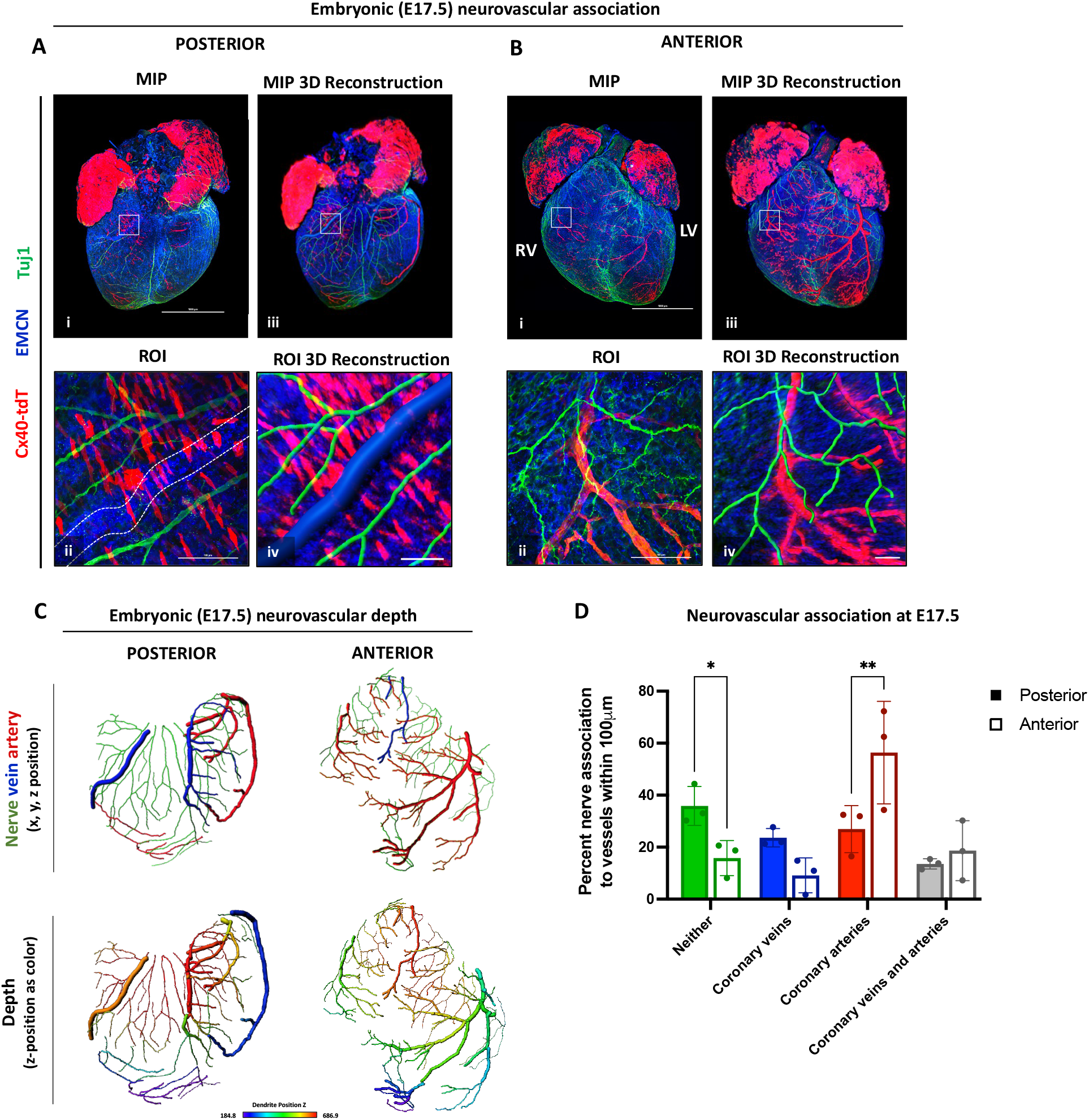
Neurovascular association shows cardiac axons align posteriorly with large veins and anteriorly with arteries. Embryonic day 17.5 (E17.5) hearts of *Cx40CreER;Rosa26^tdTomato^* mice were immunostained with Tuj1 and EMCN, followed by whole mount confocal imaging, and 3D reconstruction and modeling. The neurovascular patterning is shown as (A-Bi) a max intensity projection of the whole embryonic heart (A-B_ii_) high magnification of ROIs (A-B_iii_) 3D Imaris reconstruction of the whole embryonic heart, and (A-B_iv_) reconstruction of ROIs (n=4). (C) Neurovascular depth analysis shows that the posterior and anterior nerves are at similar depths to the vascular structure closest in proximity (n=3). (D) Quantification of percentage of nerves associated within 100um of coronary veins (blue), arteries (red) both veins and arteries (grey) or neither vessel type (green) are shown, with an increased nerve artery association in the anterior wall (n=3). Scale bars shown at 100um.

Sympathetic nerve axons innervate the heart as early as embryonic day (E) 13.5 via nerve growth factor (NGF) release by coronary veins, where sympathetic nerve axons become aligned with the veins of the posterior wall^13^. We confirmed that the axons aligned with posterior large-diameter veins from E15.5-E17.5 (**Figure 1A; Figure S1A-C**), verifying our use of EMCN to trace large-diameter veins. Since coronary arteries primarily reside on the anterior side of the ventricle at these developmental stages^38^, we hypothesized that the anterior wall would have significant nerve-artery alignment. Supporting this, as early as E16.5, axons wrapped around from the posterior wall towards arteries in the anterior wall of the heart (**Figure 1B; Figure S1D-F**). By E17.5, the nerve patterning was associated with the anterior coronary arteries (**Figures 1B_i-iv_**). These results demonstrate that nerve-artery association occurs within the anterior wall of the ventricle during late developmental stages.

We used 3D reconstruction to visualize and analyze the depth of innervation at E17.5 throughout multiple regions of the heart ventricles, where depth was defined as a function of z (**Figure 1C**). The nerves showed similar depth to their closest vascular structure, with subepicardial nerves aligning closely with the coronary veins, and myocardial nerves aligning to the coronary arteries. Interestingly, the nerves in the base of the heart were more superficial than those in the periphery and apical regions; and this patterning was consistent on both posterior and anterior walls (**Figure 1C**). Depth analysis supports the regional correlation of posterior nerve-vein alignment and anterior nerve-artery association at E17.5 (**Figure 1D**) Our results demonstrate a programmed sequential and targeted innervation pattern with respect to vessel subtype, location, and depth throughout development.

### Defining 3D Cardiac Nerve Architecture of the Postnatal Mouse Ventricle

Next, we sought to define typical nerve-vein and nerve-artery architecture in the postnatal heart. To rigorously define the 3D architecture of the postnatal heart ventricles, we considered that the spatial region of the heart is subject to natural biological variability of heart shape, size, and vascular patterning. To provide consistency between samples, we defined the architectural regions in two-fold. First, posterior and anterior sides of the heart were indicated, including right and left ventricles (LV, RV) as shown in Figure 1. Second, we used the well-defined vascular system, composed of the coronary veins and arteries, to demonstrate association of the nerves with these vessels. The coronary vessels are defined as the main right, medial and left coronary veins (RCV, MCV, LCV) and the right and left coronary arteries (RCA, LCA).

Hearts were collected at postnatal day (P) 7, a timepoint where co-maturation of the nerves and the myocardium takes place^10^. We utilized the *Cx40CreER;Rosa26^tdTomato^* mice to label coronary arteries in addition to whole mount immunostaining for nerves with Tuj1 and veins with EMCN (**Figure 2**). Whole mount imaging of the posterior wall demonstrates nerve-vein alignment, where large nerve bundles were localized near the left, medial, and right coronary veins (**Figures 2A_i-iv_**; LCV, RCV, MCV,), in agreement with previous reports^13^. Interestingly, the anterior wall demonstrates that innervation is highly localized to the right and left coronary arteries (**Figures 2B_i-iv_**; LCA, RCA). This close nerve-artery association allowed for axonal projections to directly innervate the arteries (**Figures 2B_v-vi_**). Confocal images were reconstructed in 3D by Imaris and analyzed for neurovascular association (**Figures 2C-D**). This relationship between nerves, veins, and arteries is characterized and quantified to define the typical nerve patterning of the postnatal mouse heart. The anterior wall shows a significant increase in the percent of nerves associated with the coronary arteries, in comparison to the posterior wall distribution (**Figure 2D**). Our results construct the 3D cardiac nerve architecture within the cardiovascular network in the postnatal heart, revealing a distinct innervation pattern with coronary veins and arteries in the cardiac ventricles.

**Figure 2.**
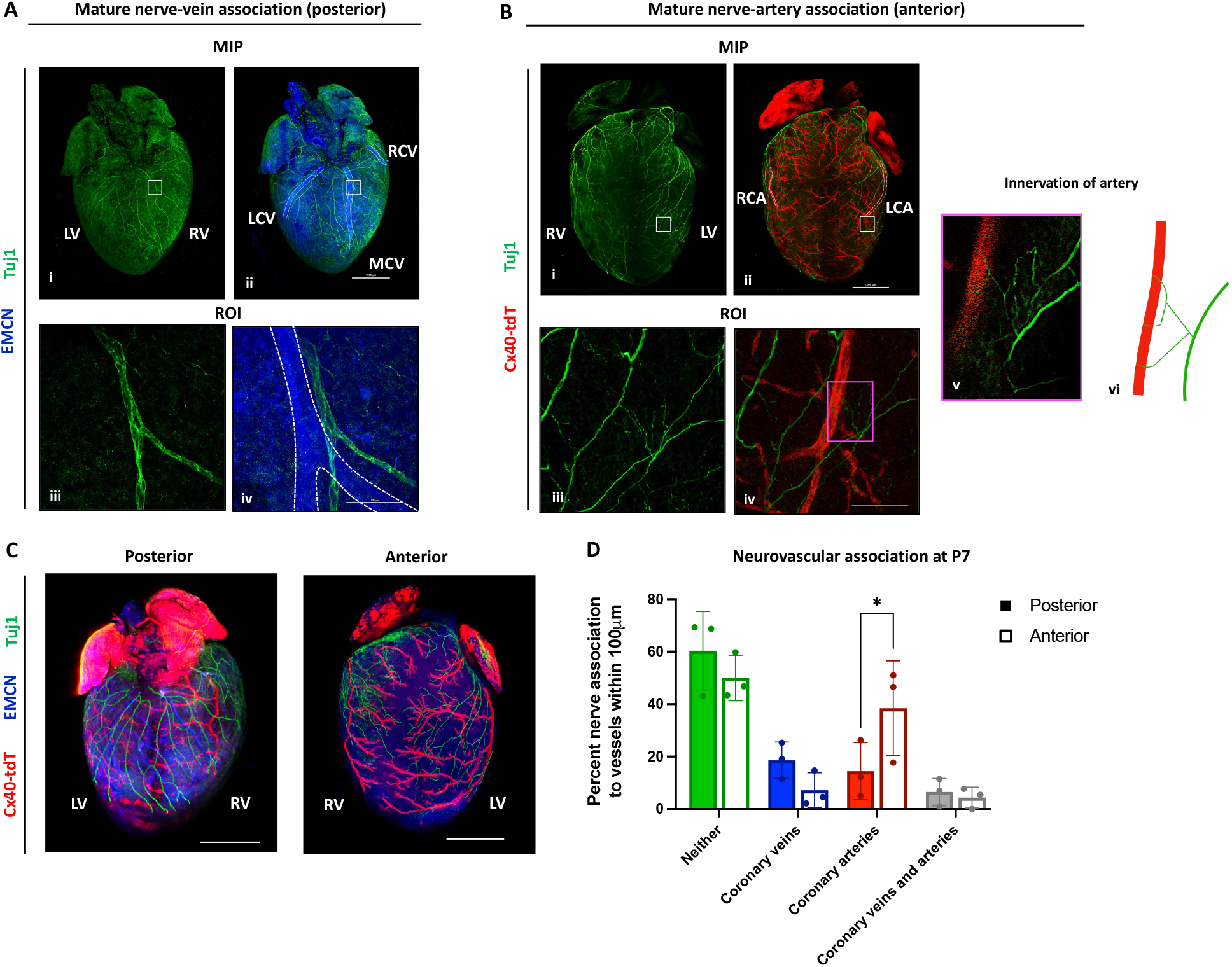
Mature 3D neurovascular architecture. P7 hearts of the *Cx40CreER;Rosa26^tdTomato^* mice were immunostained with the pan-neuronal marker Tuj1 and the endothelial cell marker endomucin (EMCN), followed by CLARITY, whole mount confocal imaging, and 3D reconstruction. (A) The posterior wall distribution of the mature cardiac nerves is shown as a max intensity projection of patterning of (A_i_) the posterior nerves alone and (A_ii_) nerve-vein association. (A_iii-iv_) The nerves align without innervating the major left, medial, or right coronary veins (LCV, MCV, RCV), shown with a representative 40x image of the region of interest (ROI) (n=4). (B) The anterior nerve architecture similarly is shown as the (B_i_) anterior nerve patterning and (B_ii_) nerve-artery association. (B_iii-vi_) Nerves align and directly innervate the right and left coronary arteries (RCA, LCA), (B_vi_) shown with a magnified z-plane of direct artery innervation (n=5). (C) Imaris 3D reconstruction highlights the neurovascular architecture of the posterior and anterior heart (n=3). (D) Quantification of percent of nerves associated within 100um of coronary veins (blue), arteries (red) both veins and arteries (grey) or neither vessel type (green) are shown, with an increased nerve artery association in the anterior wall (n=3). Scale bar is shown at 100um.

### Parasympathetic and Sympathetic Nerves Develop Synchronously and are Closely Localized Throughout Postnatal Maturation

Recent evidence suggests that parasympathetic innervation and function has an underappreciated role in the cardiac ventricles. However, the patterning and distribution of the parasympathetic innervation has not been well defined. To define the parasympathetic nerve patterning with respect to sympathetic innervation in the intact heart, we used our 3D imaging and analysis pipeline with a parasympathetic reporter mouse line and immunostaining for the sympathetic nerves. For parasympathetic nerve lineage labeling, we used the *ChATCre* knockin mouse with the *Rosa26^tdTomato^* reporter (*ChATCre;Rosa26^tdTomato^*). For sympathetic innervation labeling, we performed whole mount immunostaining with the sympathetic nerve marker Tyrosine Hydroxylase (TH). We then performed whole mount confocal imaging and 3D analysis with Imaris to construct the parasympathetic innervation of the cardiac ventricles together with sympathetic nerves (**Figure 3**).

**Figure 3.**
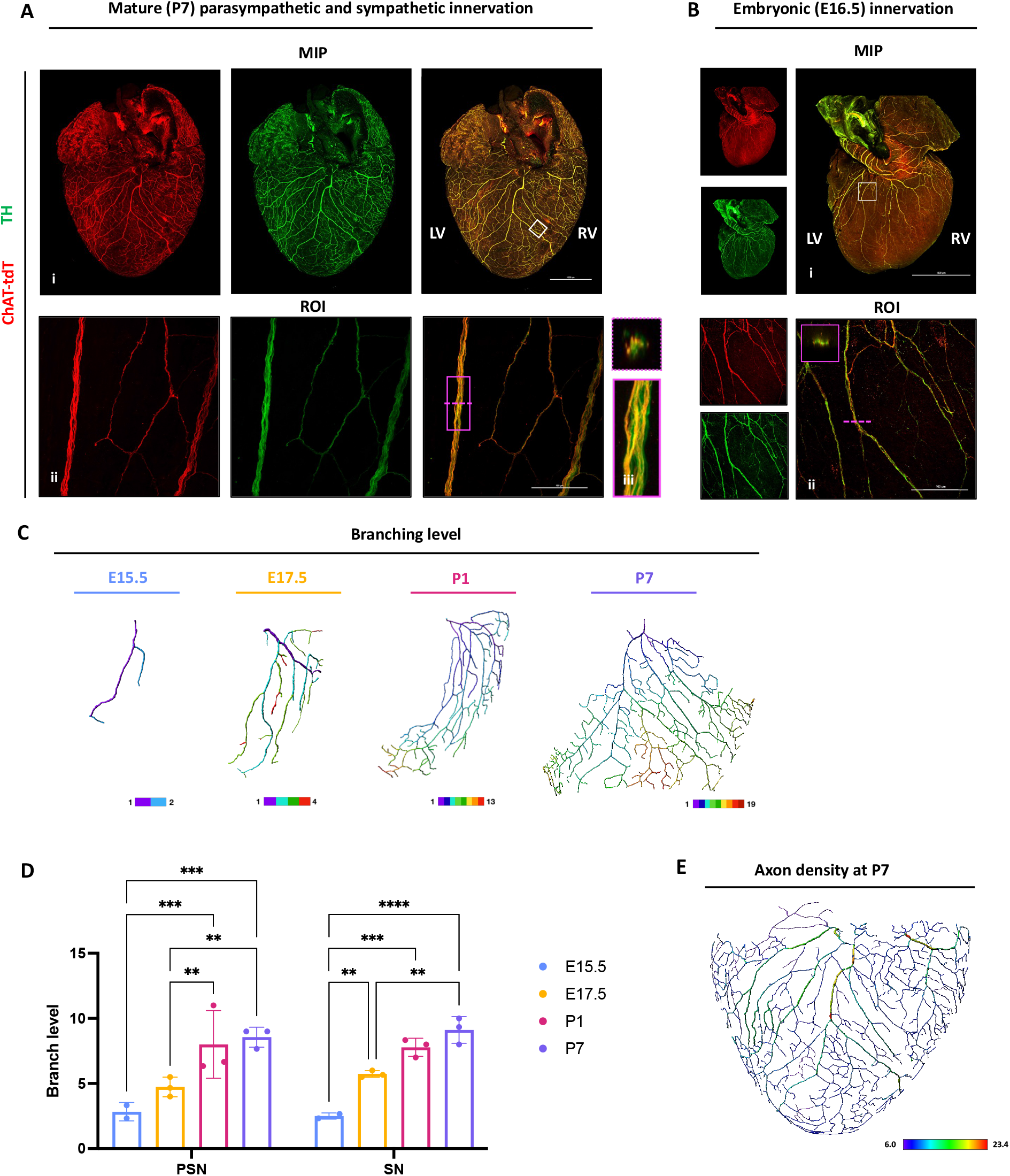
Parasympathetic and sympathetic nerves are bundled and synchronous in development. Parasympathetic reporter hearts (*ChATCre;Rosa26^tdTomato^*) were immunostained for sympathetic nerves (tyrosine hydroxylase, TH), imaged with confocal microscopy, and reconstructed with Imaris. (A) The mature nerve patterning shows (A_i_) close association between parasympathetic and sympathetic nerves, with (A_ii_) large and small nerve fibers closely aligned and the (A_iii_) large axons are bundled (n=5). (B) At E16.5, both parasympathetic and sympathetic axons extend together and are closely localized (n=6). (C-D) 3D modeling and analysis shows increased branching level during embryonic and postnatal development (n=3). (E) Density analysis of the P7 heart shows axon bundles of primary and secondary axons are at higher density than tertiary and higher branched axons.

Surprisingly, the P7 hearts of the *ChATCre;Rosa26^tdTomato^* mice showed extensive parasympathetic innervation of the ventricles (**Figures 3A_i-ii_**). Parasympathetic and sympathetic patterning was equally distributed throughout the heart. Moreover, both parasympathetic and sympathetic nerve fibers were intertwined in large bundles, as well as closely localized in smaller axon projections (**Figures 3A_ii_-A_iii_**). We then asked whether both parasympathetic and sympathetic nerve axons extend sequentially, with one guiding the other, or simultaneously, maintaining equal distribution. To investigate this, we used the *ChATCre;Rosa26^tdTomato^* mice and harvested hearts during developmental stages of axon extension at E15.5 to E17.5 (**Figure 3B, Figure S2**). At E16.5, the axons first reach the apex and demonstrate that parasympathetic and sympathetic nerves are intertwined (**Figures 3B_i-ii_**). Both parasympathetic and sympathetic nerves pattern together during earlier (E15.5) and later stages (E17.5 and P1) of heart development (**Figure S2**). No differences in distribution were seen on the posterior or anterior side of the heart (**Figure 3, Figure S3**).

To further quantify parasympathetic and sympathetic nerve patterning during embryonic and postnatal development, we used 3D reconstruction and analysis of individual branch trees, primary axons, and whole innervation networks (**Figures 3C-D, Figures S4**). Branching level, density, diameter, and patterning of parasympathetic and sympathetic nerves were investigated independently. No significant differences were found between the two, therefore we continue to describe the overall patterns identified. We analyzed branching level as a parameter of nerve development (**Figures 3C-D**). A primary axon can branch into secondary-, tertiary-, quintenary axons, and beyond, with each level assigned as branch level 1, 2, 3, respectively. The branching level increased during embryonic and early postnatal development (**Figures 3C-D**). When axons first arise around E15.5, axons have minimal branching level (approx. branch level of 2.5) and mature by P7, with a significant increase in branching level (approx. branch level of 9) (**Figure 3D**). The nerve networks also showed an increase in overall axon distances from origin, max diameter of primary filaments, and primary filament length, with a consistent trend of significance between E17.5 and P7 hearts. (**Figure S4**). Interestingly, the mature heart showed primary and secondary axons were significantly higher density compared to the tertiary branched axons and beyond (**Figure 3E, Figure S4**). These patterns identify two phases of significant nerve growth that occurs between late embryonic and early postnatal development, marked by an increase in axon branch level, distance from origin, max diameter of primary filaments, and length of primary filaments.

Our results demonstrate that parasympathetic nerves extensively innervate the cardiac ventricles and share nearly identical patterning to sympathetic nerves. This architecture is a result of synchronous parasympathetic and sympathetic axon extension, with nerve axons maintaining similar increases in branching level, distribution, and density during heart development.

### Reinnervation of the Regenerating Heart and Reestablishment of Nerve-Artery Association

Within a week after birth, the neonatal mouse heart transitions from being highly regenerative into a state of limited regenerative potential. A myocardial infarction (MI) in the non-regenerative heart causes pathological nerve remodeling, resulting in an arrythmia prone heart^26, 27^. In contrast, the neonatal heart can fully regenerate by 21 days-post MI, with full restoration of contractile and autonomic function^32^, suggesting that physiological reinnervation may take place during regeneration. Additionally, the regenerative response depends on nerve signaling^31^. Interestingly, regeneration is also dependent on collateral artery formation, where collateral arteries form at 4 days post-MI to bridge the occluded left coronary artery with the right coronary artery and mediate successful cardiac regeneration^34^. During development, artery formation precedes innervation and arterial cells recruit nerve axons to the myocardium^13, 38^. Thus, we hypothesized that newly formed collateral arteries can recruit nerve axons during cardiac regeneration.

To investigate differences in nerve remodeling in the regenerative and diseased, we performed MI surgery in regenerative (P1) or non-regenerative (P7) *Cx40CreER;Rosa26^tdTomato^* mice and collected hearts at 7- or 21 days-post MI. Hearts underwent tissue clearing (**Figure 4A**), followed by whole mount immunostaining with the pan-neuronal marker Tuj1 (**Figure 4**). We first investigated innervation remodeling in the non-regenerated heart (**Figure 4B**). We demonstrate that the non-regenerated heart is starkly denervated in the infarct zone at 21 days post-MI (**Figure 4B**). This is similar to the denervation seen in the infarcted adult human heart^39^ and interestingly, the degree of denervation, rather than infarction size, is an accurate predictor of ventricular arrhythmias^40^. We then investigated how this compared to nerve remodeling in the regenerative heart. Since collateral arteries are formed by 4 days post-MI in the regenerating heart^34^, we began by investigating nerve remodeling shortly after at 7 days post-MI (**Figure 4C**). Interestingly, whole mount imaging demonstrates that although collateral arteries start to appear in the infarct zone at P4, the infarct zone remains denervated at P7 (**Figure 4C**). We then explored the remodeling of nerve-artery architecture in the fully regenerated myocardium at 21 days post-MI (**Figures 4D)**. Our results demonstrate extensive reinnervation of the regenerated myocardium (**Figure 4D_i_**). Excitingly, the nerves show reestablishment of nerve-artery connection with the collateral arteries, suggesting targeted innervation takes place (**Figure 4D_ii_**).

**Figure 4.**
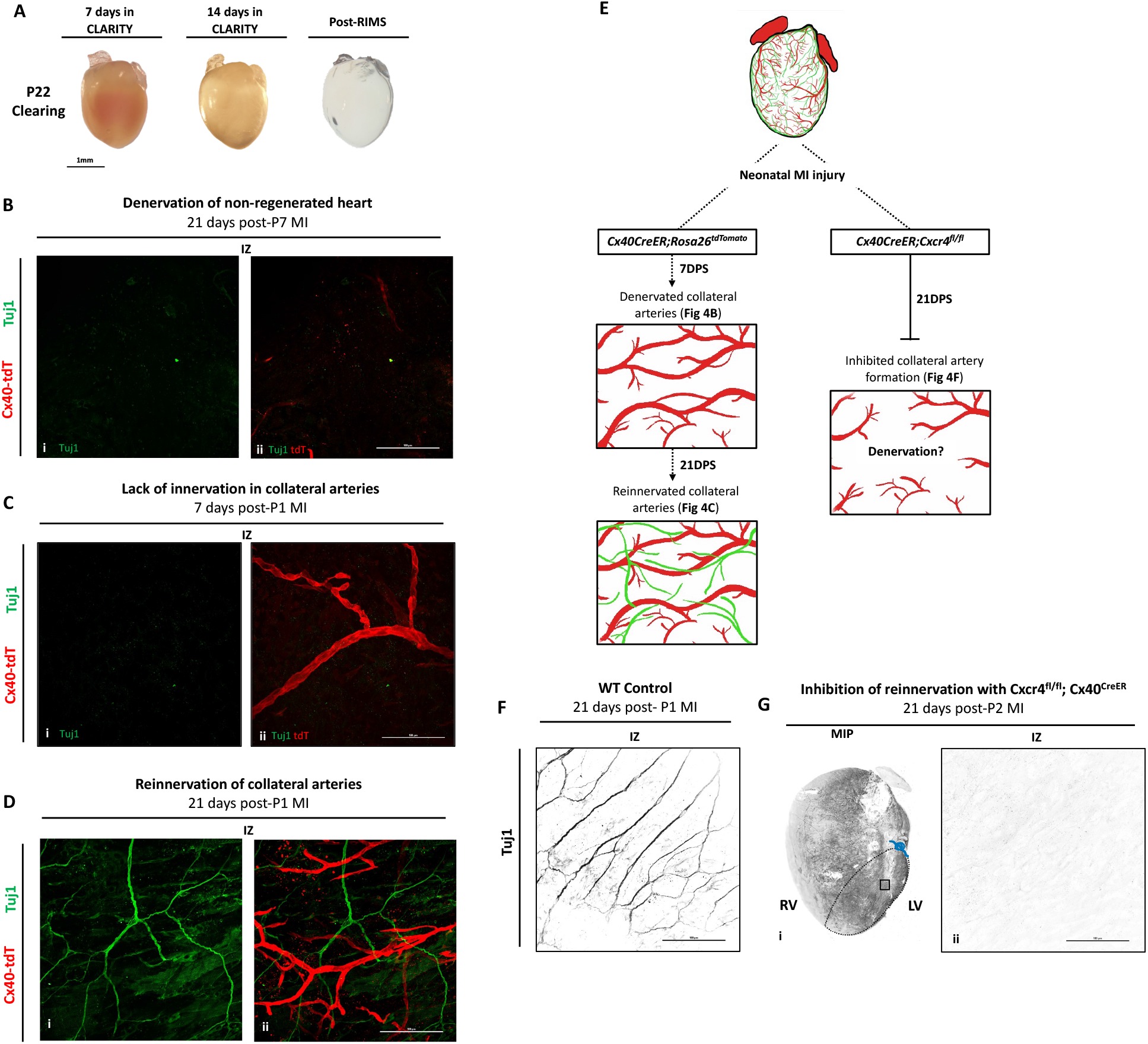
Reinnervation of the regenerating heart is dependent on collateral artery formation. MI was performed in *Cx40CreER;Rosa26^tdTomato^* mice during the regenerative window (P1 or P2) and hearts were collected at 7- or 21-days post-MI. (A) Hearts underwent tissue clearing and nerve immunostaining with Tuj1. (Bi-ii) 7 days after P1 MI, (Bi) the infarct zone (IZ) is denervated, (Bii) including the newly formed collateral arteries (n=6). (Ci-ii) At 21 days after P1 MI, (Ci) the regenerated tissue is reinnervated, (Cii) with nerves showing artery association (n=4). (D) To determine whether reinnervation during regeneration is dependent on collateral artery formation, we used the *Cx40CreER;Cxcr4^fl/fl^* mouse to inhibit migration of arterial cells post-MI. (Fi-ii) At 21 days post-MI, Tuj1 immunostaining showed the IZ remained denervated in the *Cx40CreER;Cxcr4^fl/fl^* mice (n=4), in comparison to (E) control regenerating hearts (n=4). Black dashed border indicates the IZ and the blue represents the suture site.

To determine whether reinnervation is dependent on collateral artery formation, we impaired collateral artery formation using the (*Cx40CreER;Cxcr4^fl/fl^* mouse^34^ (**Figures 4E-G**). Cxcr4 is a chemokine receptor expressed in arterial endothelial cells that responds to the chemotactic ligand Cxcl12^41–43^. The Cxcl12/Cxcr4 axis is important for arterial cell migration and artery formation during development and regeneration, where deletion of *Cxcr*4 using the *Cx40CreER* mice impairs collateral artery formation during neonatal heart regeneration^34^ We injected *Cx40CreER;Cxcr4^fl/fl^* pups at P0 with a single dose of tamoxifen and MI was performed within the regenerative window at P2 (**Figure 4E**). Hearts were collected at 21 days-post-MI and underwent whole-mount clearing and immunostaining for Tuj1. Strikingly, the infarcted area was denervated, in stark contrast to the regenerating wild type controls (**Figure 4F-G**), demonstrating that reinnervation is dependent on collateral artery formation. This is the first evidence of targeted reinnervation in the regenerating myocardium, and that this reinnervation is dependent on collateral artery formation.

### Physiological Reinnervation of the Regenerated Myocardium

Nerve remodeling following cardiac injury in the adult mammalian heart is a prominent hallmark of the pathology of heart failure. In contrast, following a neonatal mouse heart injury, the heart regenerates normally, and our results demonstrate reinnervation of the regenerating myocardium. This suggests that reinnervation patterns following adult injury and during neonatal heart regeneration vary widely, and proper innervation patterning is crucial for survival and restoring normal autonomic function of the heart.

To distinguish between nerve remodeling in regenerating and non-regenerating hearts, we performed MI surgery in *ChATCre; Rosa26^tdTomato^* mice at the regenerative (P1) and non-regenerative (P7) timepoints and collected hearts at 21 days post-MI. Collected hearts were further stained with tdTomato and TH to identify both parasympathetic and sympathetic nerve patterning, respectively. The control uninjured hearts at P22 demonstrate both parasympathetic and sympathetic nerve bundling, and patterning as expected (**Figure 5A**). In contrast, the infarct zone of the P7 non-regenerating heart at 21 days post-MI showed complete denervation (**Figure 5B_iii_**). Furthermore, the non-regenerating hearts demonstrate sympathetic nerve hyperinnervation at the border zone, a distinctive feature of pathological nerve remodeling (**Figures 5B_ii_**)^29^. Remarkably, the regenerating myocardium at 21 days post-MI of the P1 heart show restoration of both parasympathetic and sympathetic nerve architecture (**Figures 5C_i-iii_**), and the border zone shows large axon bundles of parasympathetic and sympathetic nerves (**Figure 5C_ii_**). This reinnervation of the parasympathetic and sympathetic nerves covers the newly regenerated myocardium (**Figure 5C_iii_**). Collectively, the reestablishment of parasympathetic and sympathetic nerves (**Figure 5C**) along with direct artery reinnervation (**Figure 4C**) demonstrates that physiological reinnervation takes place during regeneration. This is in stark contrast to the non-regenerating heart, which shows denervation of the infarct zone and pathological reinnervation, including sympathetic hyperinnervation at the border zone.

**Figure 5.**
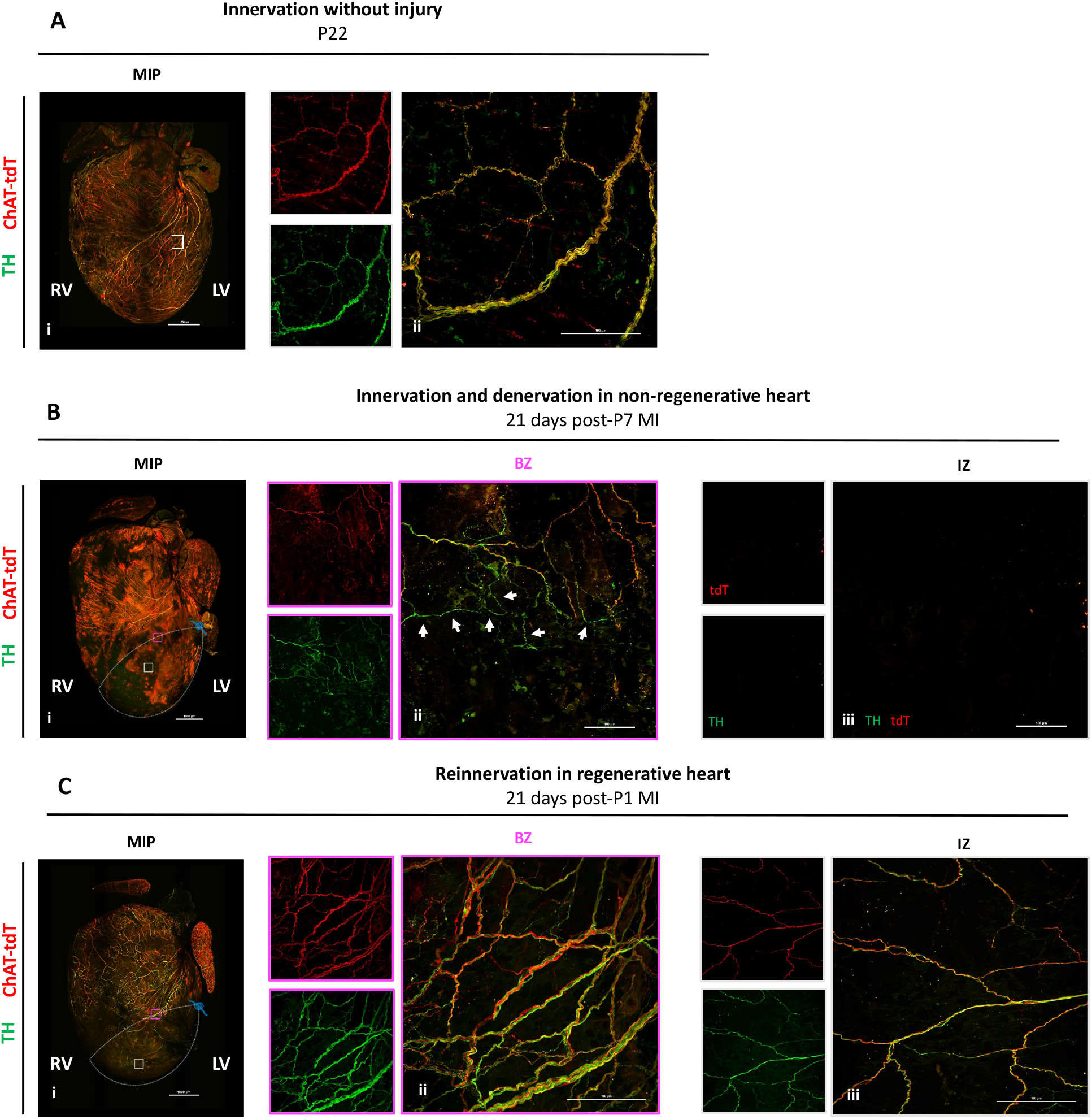
Parasympathetic and sympathetic nerves reinnervate the regenerated myocardium. Myocardial Infarction (MI) was performed on *ChATCre;Rosa26^tdTomato^* mice at regenerative (P1) and non-regenerative (P7) timepoints. Hearts collected at 21 days post-MI, and immunostained for tyrosine hydroxylase (TH) and imaged with confocal microscopy. (A_i=ii_) Uninjured hearts in adult (P22) mice show parasympathetic and sympathetic nerve entanglement in (A_i_) whole mount and (A_ii_) in the medial region of the myocardium (n=6). (B_i=iii_) Non-regenerating hearts show pathological remodeling of the nerves. (B_ii_) The border zone (BZ) shows reinnervation of sympathetic axons, independent of parasympathetic axons. (B_iii_) The infarct zone (IZ) is denervated (n=6). (C_i=iii_) The regenerated heart shows reestablished parasympathetic and sympathetic axon bundling. (C_ii_) The BZ shows large nerve bundles of both nerve subtypes. (C_iii_) The IZ shows reinnervation of parasympathetic and sympathetic axons (n=5). White dashed border indicates the IZ and the blue represents the suture site.

## DISCUSSION

Neural regulation of the cardiovascular system has been recognized for a long time, however; the importance of neurocardiology in cardiovascular health and disease is beginning to be appreciated. Recent studies aimed at identifying the cellular and molecular makeup of the intrinsic cardiac nervous system and the sinoatrial node underscore the importance of elucidating the innervation patterns and function of different parts of the cardiac nervous system^4, 5^. Sympathetic innervation of cardiac ventricles has been studied extensively^6, 13^, however; parasympathetic innervation has been underappreciated due to technical limitations^16^. In our experience, whole mount immunolabeling of ChAT is technically challenging. Furthermore, 2D analysis from histological sections is inaccurate and cannot reconstruct the complex patterns of nerves and networks with other cell types. Furthermore, endogenous labeling using a lineage reporter alone is prone to quenching (**Figure S5**). To overcome this limitation, we utilized the *ChATCre; Rosa26^tdTomato^* model together with whole mount immunostaining for tdTomato and tissue clearing, which allowed for an accurate view of the parasympathetic nervous system. Our finding of extensive parasympathetic innervation of the ventricles indicates potential intracellular dynamics and physiological influences between parasympathetic nerves and the surrounding heart tissue. Furthermore, we demonstrate a spatiotemporal innervation pattern of cardiac ventricles with respect to coronary veins and arteries.

Cardiac injury results in neuronal degeneration and pathological nerve remodeling, which leads to a disruption in heart innervation patterns^26, 27, 29^. This neural remodeling and pathological innervation that occurs following injury leads to fatal arrhythmias^25^. Nerves have been therapeutically targeted using neuromodulation approaches, such as vagal nerve stimulation and sympathetic nerve denervation^44–47^. Some studies demonstrate promising outcomes, however; our lack of understanding of cardiac nerve development and remodeling hampers our understanding of the mechanisms by which these approaches modulate reinnervation and function^48^.

Neonatal mice can regenerate their hearts following injury for a brief window after birth^32^. Cardiac nerves have been demonstrated to regulate cardiomyocyte proliferation and neonatal heart regeneration^31^. Furthermore, autonomic heart functions are restored following regeneration, suggesting that physiological reinnervation take place, which contrasts with the pathological innervation that takes place following adult cardiac injury. Additionally, from a clinical perspective, it has been well established that heart transplant recipients receive a completely de-innervated heart, which eventually becomes partially innervated by the host^49^. Thus, understanding the development and plasticity of cardiac innervation is a unique approach to stimulate physiological innervation and cardiac regeneration.

Remarkably, we demonstrate for the first time the reestablishment of physiological innervation of the regenerating myocardium, suggesting that this reinnervation likely preserves autonomic function of the regenerated myocardium in contrast to the non-regenerating hearts. Furthermore, we demonstrate that physiological innervation is dependent on collateral artery formation, where inhibition of collateral arteries following injury blocks this reinnervation. These results suggest that promoting collateral artery formation can be targeted to promote both physiological reinnervation and cardiac regeneration following injury.

Our study reveals new insights into cardiac innervation during development, disease, and regeneration. However, it remains unclear whether a distinct gene regulatory network mediates physiological and pathological innervation. Furthermore, identifying the signals by which coronary arteries regulate reinnervation during regeneration can play an important role in treatment of autonomic dysfunction, as well as promote cardiac repair following injury. Our results provide a framework to start dissecting the cellular and molecular networks that guide cardiac innervation and regeneration.

## Acknowledgments

Funding for this project was provided by an AHA Predoctoral Fellowship 829586 (R.J.S.), NIH/NHLBI R56 HL155617 (A.I.M.), NIH/NHLBI R01 HL166256 (A.I.M.), DOD W81XWH2210094 (A.I.M.), and NIH/NIGMS R35 GM 134865 (A.A.). We thank Lucile Miquerol and Kristy Red-Horse for generously sharing the *Cx40CreER* mice. We thank Lance Rodenkirch and the Optical Imaging Core (Grant 1S10OD025040-01) for imaging support.

