## Supplemental Figures for "Defining Cardiac Nerve Architecture During Development, Disease, and Regeneration": BioRxiv Supplementary Figures.pdf

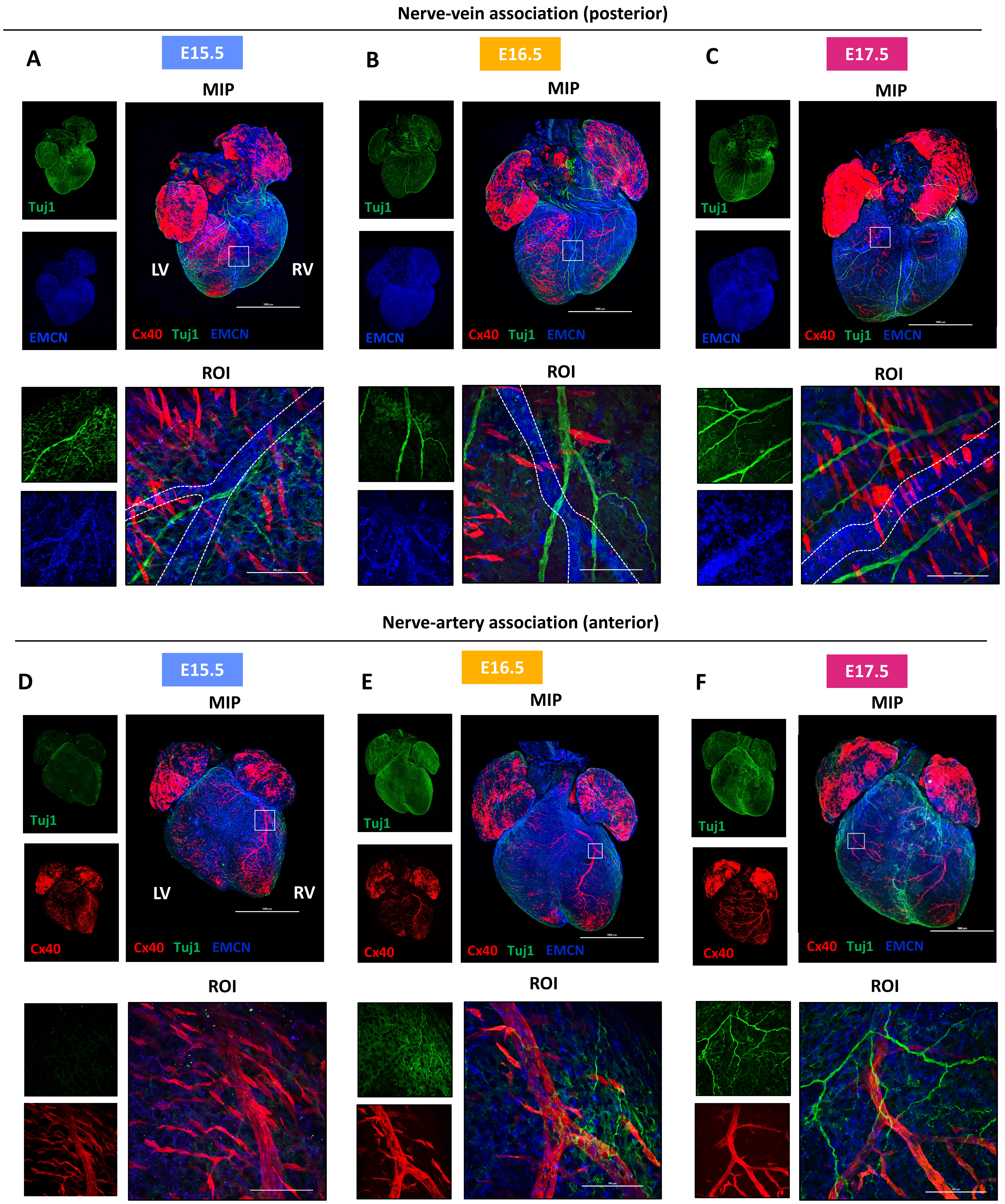


**Figure S1. Embryonic nerve-vein and nerve-artery association.** Embryonic hearts were immunostained for nerves (Tuj1), veins (EMCN) and arteries (Cx40-tdTomato) and imaged using confocal microscopy. The embryonic nerve development shows axon extension occurs from E15.5-E17.5. (A-C) The nerves first appear on the posterior heart at E15.5 and show association with the veins at (A) E15.5, (B) 16.5) and (C) 17.5. (D-F) The anterior portion of the heart the begins to be innervated at E16.5 and axons associate with the coronary arteries by E17.5.

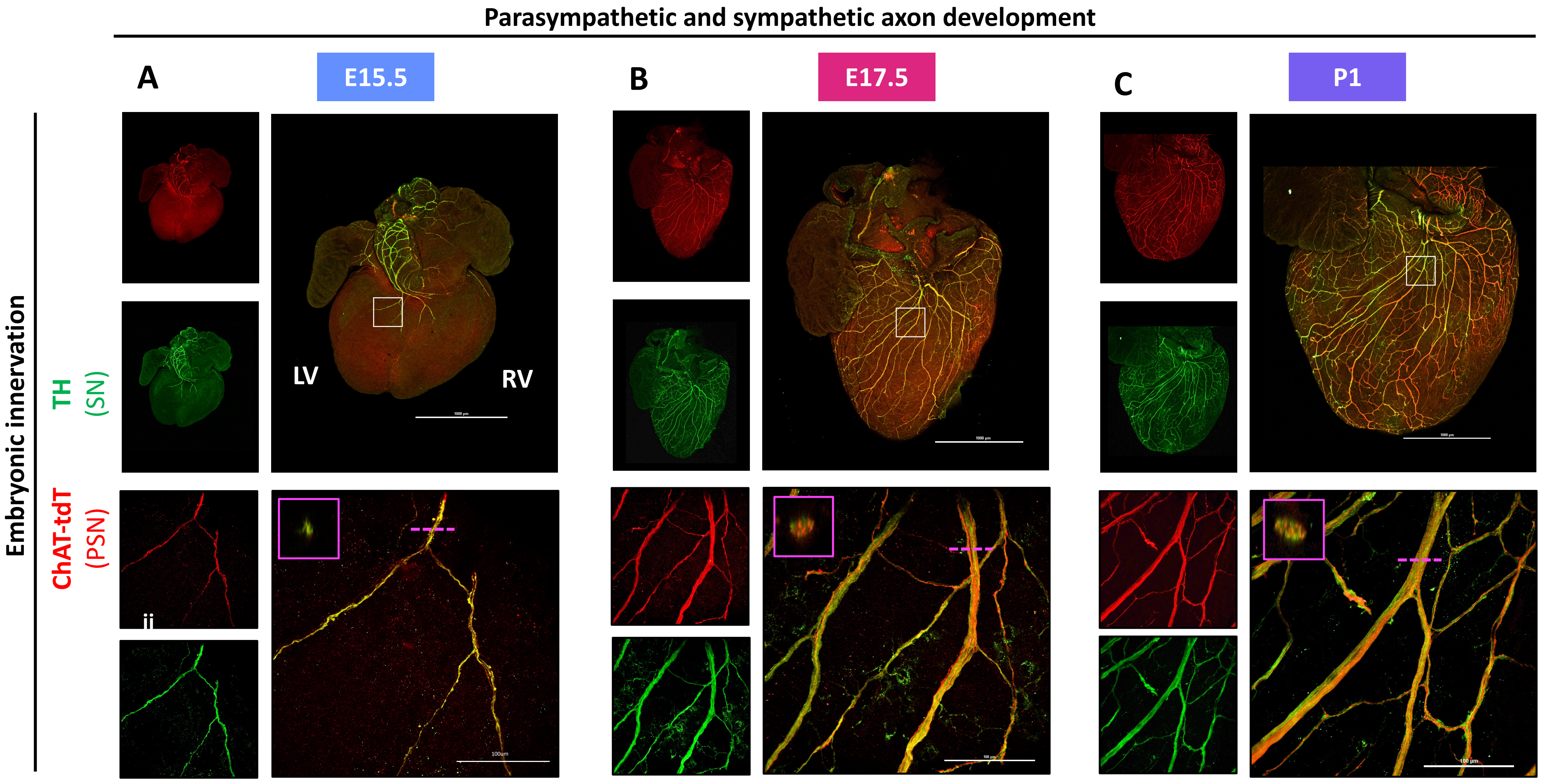


**Figure S2. Embryonic development of parasympathetic and sympathetic nerves.** Parasympathetic reporter hearts (*ChATCre;Rosa26*$^{tdTomato}$) were harvested during embryonic development. (A) At E15.5 nerves first appear in the ventricles, with PSNs and SNs showing close association and bundling (n=7); this trend continues throughout (B) late embryonic development, E17.7 (n=6), and (C) early postnatal development, P1 (n=5)

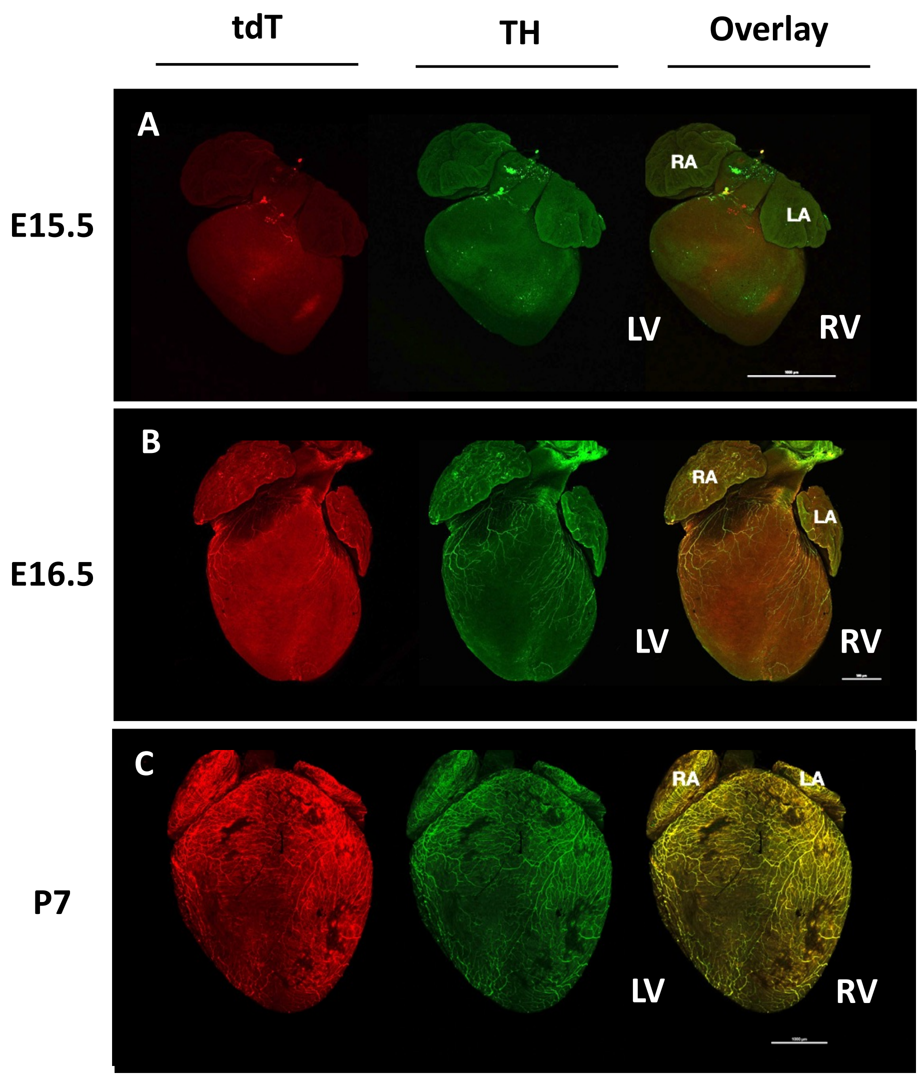


**Figure S3. Posterior images of axon development.** The PSNs and SNs in the anterior heart are (A) not present at E15.5, (B) begin to innervate synchronously at E16.5 and (C) are densely and closely associated at P7. Right and left atria and ventricles are defined (RA, LA, RV, LV)

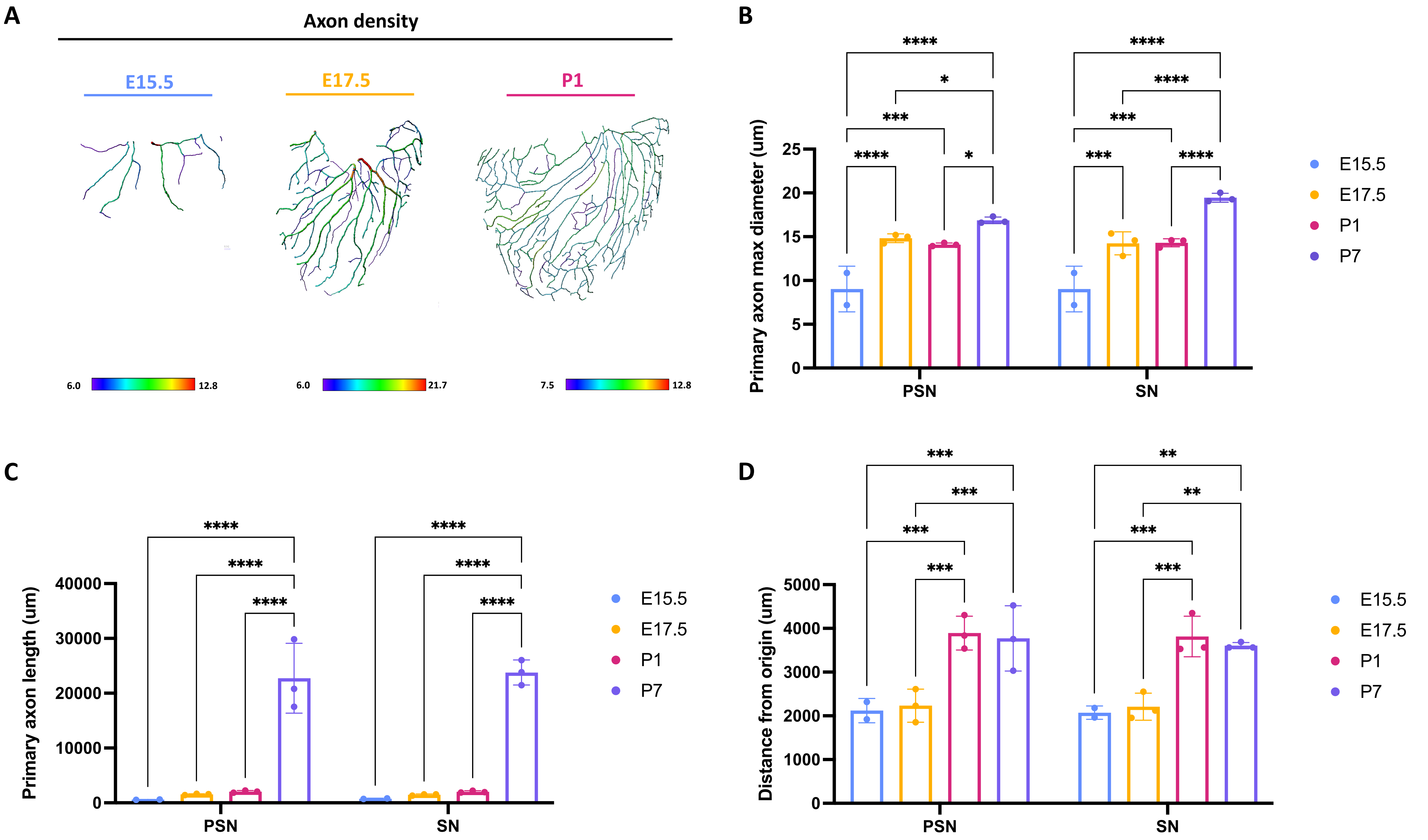


**Figure S4. Analysis of embryonic and postnatal axon development.** (A) Density distribution in developing nerve networks. Axon density is shown at E15.5, E17.5 and P1 for the entire parasympathetic nerve networks. Quantification of (B) primary axon max diameter (C) primary axon length and (D) distance from origin for parasympathetic (PSN) and sympathetic (SN) nerves during embryonic and postnatal development.

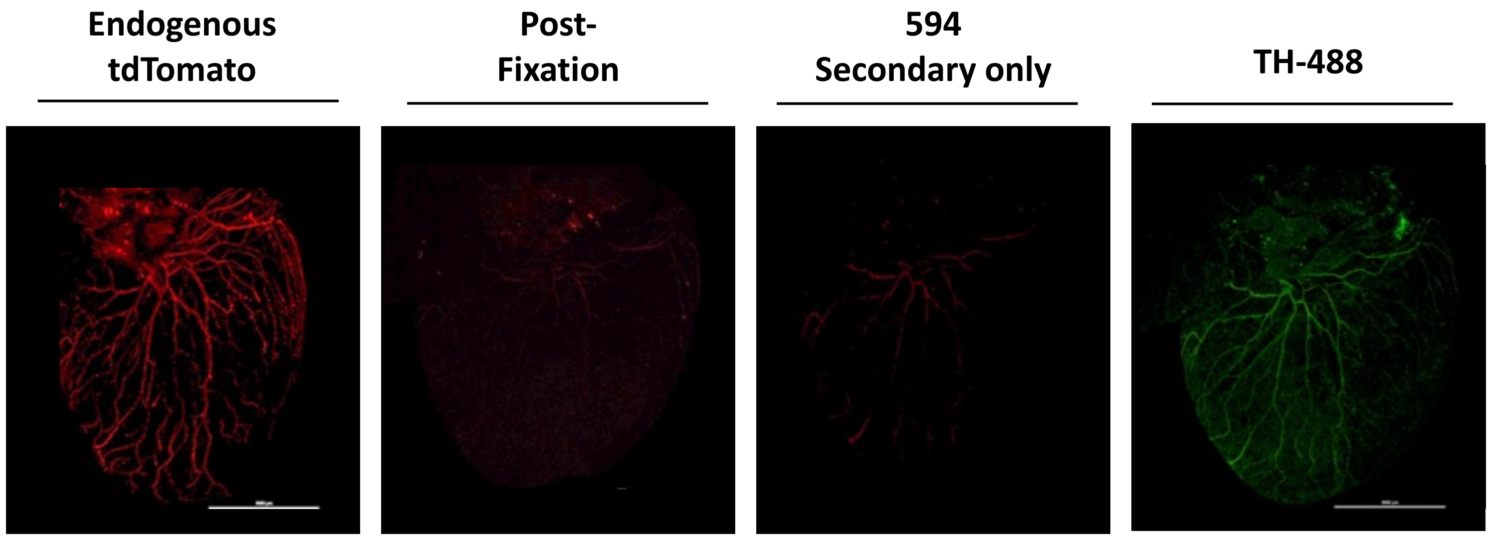


**Figure S5. ChAT-Tdtomato signal is quenched post-fixation.** ChAT-Tdtomato staining control to show secondary is specifically binding to tdTomato, shown by whole-mount of P1 hearts. (A) Endogenous TdTomato expression is fluorescent. (B) Post-fixation in PFA, this endogenous signal is quenched. (C)Adding in 594 secondary only without tdTomato primary produces no off-target binding. (D) Staining with Tyrosine Hydroxylase (TH) primary and matching 488 secondary shows the 488 specific to the TH epitope binds specifically.
